# Autofluorescence lifetime flow cytometry with time‐correlated single photon counting

**DOI:** 10.1101/2024.05.15.594394

**Authors:** Kayvan Samimi, Ojaswi Pasachhe, Emmanuel Contreras Guzman, Jeremiah Riendeau, Amani A. Gillette, Dan L. Pham, Kasia J. Wiech, Darcie L. Moore, Melissa C. Skala

**Affiliations:** Morgridge Institute for Research, Madison, WI, USA; Department of Biomedical Engineering, University of Wisconsin, Madison, WI, USA; Department of Neuroscience, University of Wisconsin, Madison, WI, USA

**Keywords:** flow cytometry, label‐free sensing, fluorescence lifetime, metabolism, NAD(P)H, time tagger, single‐cell analysis

## Abstract

Autofluorescence lifetime imaging microscopy (FLIM) is sensitive to metabolic changes in single cells based on changes in the protein-binding activities of the metabolic co-enzymes NAD(P)H. However, FLIM typically relies on time-correlated single-photon counting (TCSPC) detection electronics on laser-scanning microscopes, which are expensive, low-throughput, and require substantial post-processing time for cell segmentation and analysis. Here, we present a fluorescence lifetime-sensitive flow cytometer that offers the same TCSPC temporal resolution in a flow geometry, with low-cost single-photon excitation sources, a throughput of tens of cells per second, and real-time single-cell analysis. The system uses a 375nm picosecond-pulsed diode laser operating at 50MHz, alkali photomultiplier tubes, an FPGA-based time tagger, and can provide real-time phasor-based classification (*i.e*., gating) of flowing cells. A CMOS camera produces simultaneous brightfield images using far-red illumination. A second PMT provides two-color analysis. Cells are injected into the microfluidic channel using a syringe pump at 2-5 mm/s with nearly 5ms integration time per cell, resulting in a light dose of 2.65 J/cm^2^ that is well below damage thresholds (25 J/cm^2^ at 375 nm). Our results show that cells remain viable after measurement, and the system is sensitive to autofluorescence lifetime changes in Jurkat T cells with metabolic perturbation (sodium cyanide), quiescent vs. activated (CD3/CD28/CD2) primary human T cells, and quiescent vs. activated primary adult mouse neural stem cells, consistent with prior studies using multiphoton FLIM. This TCSPC-based autofluorescence lifetime flow cytometer provides a valuable label-free method for real-time analysis of single-cell function and metabolism with higher throughput than laser-scanning microscopy systems.

## 1. Introduction

Fluorescence lifetime imaging microscopy (FLIM) of metabolic co-factors nicotinamide adenine dinucleotide (NADH), nicotinamide adenine dinucleotide phosphate (NADPH), and flavin adenine dinucleotide (FAD) is often used to monitor the metabolism of single cells [1–5]. In their reduced form, NADH and NADPH have an absorption peak at 350 nm and emission peak at 460 nm with indistinguishable fluorescence properties, and their combined emission is denoted by NAD(P)H [6]. In its oxidized form, FAD has absorption peaks at 355 nm and 450 nm, and an emission peak at 530 nm [6]. The fluorescence lifetime of NAD(P)H is shorter in its free state (∼400 ps) compared to its protein-bound state (∼1 to 5 ns) [6,7]. Conversely, FAD has longer fluorescence lifetime in its free state (2.3 to 2.9 ns) compared to its protein-bound state (<0.1 ns) [6,7]. Previous studies have used these autofluorescence lifetimes as biomarkers for disease states that are associated with metabolic dysfunction, including cancer [8–11], autoimmune diseases [12], cardiovascular diseases [13,14], and diabetes [15,16]. Importantly, autofluorescence lifetime measurements provide rapid insight into molecular interactions and cellular metabolism without the use of exogenous reagents. As such, this technique can be readily applied to study novel biological and biopharmaceutical questions, without the upfront need to develop cell-specific labels and markers that can be costly, labor-intensive, have poor reproducibility, or be prohibited in some applications (*e.g*., T cell or stem cell therapies in patients).

The low absorption coefficient and low quantum yield of endogenous fluorophores typically requires laser-scanning multiphoton or confocal microscopes performing time-correlated single-photon counting (TCSPC) to provide sufficient signal-to-noise and signal-to-background ratios for successful analysis of complex decay profiles [17]. The image acquisition time in these FLIM systems is typically on the order of a minute per field of view. The slow acquisition speed and the microscopy geometry make scanning TCSPC FLIM unsuitable for rapid assays involving large numbers of cells, or when cell sorting is desired. Therefore, there has been a continued interest in performing these label-free autofluorescence lifetime measurements in a flow geometry [18–22].

Previous flow studies have used frequency-domain techniques in which the excitation laser is amplitude modulated and the NAD(P)H lifetime is estimated from the emission demodulation and phase shift relative to the excitation laser [18–20]. While the frequency-domain technique is readily compatible with existing flow cytometers in terms of laser power (tens of milliwatts) and flow velocity (tens of microseconds cell transit times), it is best suited to bright fluorophores that produce a modulated emission with high signal-to-noise ratio from which the phase shift (and subsequently the fluorescence lifetime) can be reliably estimated [23]. In contrast, endogenous fluorophores like NAD(P)H typically have low absorption coefficients and quantum yields [6,24,25]. Additionally, the short lifetimes of these endogenous fluorophores (*e.g*., 0.4 ns for free NAD(P)H) require large modulation frequencies (tens of MHz) to produce large phase shifts, which in turn requires fast (multi-GHz) digitizers for recording and frequency analysis of the modulated emissions. Therefore, the frequency-domain technique (with a single modulation and demodulation frequency) is not the most photon-efficient technique for measuring the fluorescence lifetime of endogenous fluorophores. In contrast, the TCSPC technique allows for efficient recovery of the time-resolved decay profile along with fitting to resolve multi-exponential decay components and their lifetimes [26].

Here, we present a single-photon excited lifetime-sensitive flow cytometer that acquires whole-cell TCSPC NAD(P)H fluorescence decays from single cells in a microfluidic flow channel at rates up to 100 cells per second and provides real-time phasor-based classification (*i.e*., gating) of the cells. This flow geometry is attractive for interrogating multiple biological questions including immune cell activation, stem cell differentiation, optimization of cell therapy manufacturing, rapid drug screening, and other situations in which touch-free metabolic assessment of cells is desirable. Importantly, as labeled flow-based analyses are common in many fields, this tool provides seamless integration into existing operational procedures without the need for additional training or deep understanding of new FLIM instruments or concepts.

## 2. Materials and Methods

### 2.1 Instrumentation

A lifetime-sensitive microfluidic flow cytometer with two color channels was built around an inverted Ti-S microscope (Nikon Instruments Inc., Melville, NY) equipped with a motorized PZ-2000 stage (Applied Scientific Instrumentation, Eugene, OR). The schematic diagram of the system is shown in Fig. 1. For autofluorescence excitation, we used a QuixX 375-70PS picosecond-pulsed diode laser (Omicron-Laserage Laserprodukte GmbH, Rodgau, Germany) operated at 50 MHz pulse repetition rate (*i.e*., 20 ns interval) and producing narrow (<90 ps width) pulses. The 375 nm laser beam (Ø1 mm 1/e^2^) was coupled through the epi-illumination port, tube lens (f = 165 mm), 405 nm long-pass (LP) dichroic mirror, and a 100× 1.3 NA oil-immersion objective lens (Nikon) to produce a Ø12 µm full width at half maximum (FWHM) Gaussian profile at the focal plane (Fig. 1B) with an average power of 0.6 mW. This spot size was chosen to closely match the size of a single cell and minimize background excitation to produce optimal signal to background ratio. For the beads experiment a neutral-density (ND) filter with optical density of 3 was used to lower the excitation laser power.

**Fig. 1:**
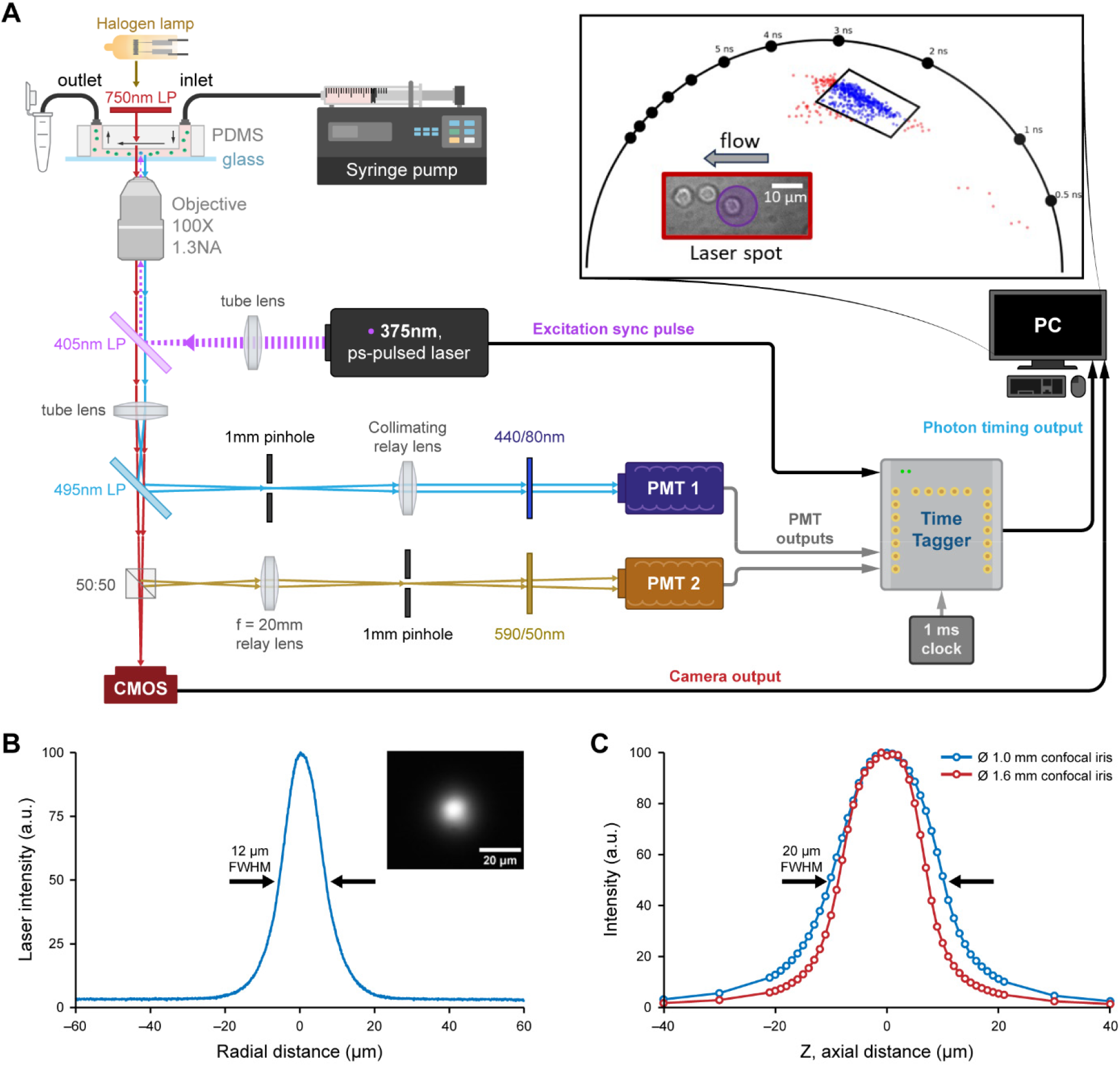
Schematic diagram of the autofluorescence lifetime flow cytometer. (A) The system includes a 375 nm picosecond-pulsed diode laser operated at 50 MHz pulse repetition rate, two ultra bialkali photon-counting PMTs, an FPGA-based time tagger connected to a computer via USB 3.0 connection, a CMOS camera with near-infrared illumination for bright field view of the flow cell. The microfluidic flow cell is made from PDMS bonded to No. 1 coverglass, and the sample cells are injected through the channel using a high-precision syringe pump. Autofluorescence decays from single cells are recorded and their phasor representation is visualized on the universal semi-circle in real time. A user-defined polygon gate on the phasor plot enables real-time detection of cells (blue vs. red dots) from a subset of interest. (B) The excitation laser beam at the focal plane has a Ø 12 µm full width at half maximum Gaussian profile. Inset shows the CMOS view of the excitation beam illuminating a thin film of saturated coumarin-6 solution in ethanol. (C) The axial profile of the confocal volume has a measured 20 µm full width at half maximum using a Ø 1 mm confocal iris.

NAD(P)H autofluorescence emissions were collected by the same objective lens and detected using an ultra bialkali H10721P-210 PMT (Hamamatsu Corp., Bridgewater, NJ) placed on a side port after a 495nm LP dichroic mirror, a Ø1 mm confocal iris, and a 440/80nm emission filter (Semrock Inc., Rochester, NY). A second PMT collected longer wavelength (590/50nm) signal from autofluorescence or exogenous labels when applicable. The confocal iris had a back-projected diameter of 10 µm at the sample focal plane, which matches the lateral size of the illumination spot and single cells. The corresponding axial profile of the confocal volume had a measured 20 µm FWHM (Fig. 1C).

Time-resolved measurement of the autofluorescence emissions was achieved by time-correlated single photon counting (TCSPC). A field programmable gate array (FPGA)-based time tagger device, Time Tagger Ultra (Swabian Instruments GmbH, Stuttgart, Germany) was used to measure the time difference between an excitation laser pulse and the detection of an autofluorescence photon by a PMT. These photon time tags were continuously transferred via USB 3.0 to a host computer that compiled them into autofluorescence decay histograms over many cycles of the excitation laser. An external function generator supplied a 1 kHz clock signal that defined the collection period of successive histograms.

A acA1300-200 um CMOS camera (Basler Inc., Exton, PA) with near-infrared illumination from the halogen lamp with 750nm LP Schott glass filter (Thorlabs Inc., Newton, NJ) provided brightfield images of the flow cell to focus the optics and verify proper flow through the channel. This near-infrared illumination does not interfere with the autofluorescence spectrum of NAD(P)H.

### 2.2 Microfluidic flow cell

The flow cells were made from polydimethylsiloxane (PDMS), SYLGARD 184 (Dow Inc., Midland, MI) for rapid prototyping. Negative molds of the microfluidic channel were designed in SolidWorks (Dassault Systèmes Corp., Paris, France) and fabricated using a resin 3D printer, Viper SI2 (3D Systems Inc., Rock Hill, SC). PDMS was cured in the 3D-printed molds at 90°C for 30 minutes. Next, it was demolded, plasma cleaned (PDC-001, Harrick Plasma Inc., Ithaca, NY), and bonded to No.1 cover glass that formed the bottom surface of the channel and the optical window for the 100x objective lens. A biopsy punch to the top of the PDMS channel provided openings for inlet and outlet tubing connections. Samples were injected through the inlet using a 100 µL glass syringe (Hamilton Company, Reno, NV) and a syringe pump (Pump 11 Elite, Harvard Apparatus, Holliston, MA).

Several flow cell designs with different focusing sheath fluid configurations were implemented. However, except for the beads flow experiments (Fig. 3), all data presented in this manuscript was acquired using a simple flow channel (as in Fig. 1) with a 50-µm square profile, one inlet and one outlet, and no focusing sheath fluid. For the beads flow experiment, a flow channel with both horizontal and vertical focusing sheath inlets, producing a 25-µm core was used.

The flow velocity determines the transit time of the cells through the Ø12 µm laser illumination spot. Given the empirical autofluorescence photon count rate of 2 million photons per second observed from T cells with this setup, and a target photon count of 10,000 photons per cell for accurate two-component lifetime estimation [27,28], an integration time of 5 ms per cell was required. Therefore, a flow velocity of 2.4 mm/s (*i.e*., 12µm/5ms) or slower was necessary. At this velocity, the ultraviolet light dose from the 0.6 mW laser beam onto cells crossing the Ø12 µm illumination spot is 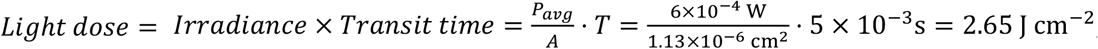, which is about 10x below the viable light dose at this wavelength [29].

### 2.3 Custom TCSPC algorithm for cell transit detection

A custom TCSPC algorithm was programmed in Python using the Time Tagger application programming interface (API) [30] to compile autofluorescence decay histograms from the stream of photon detection time tags and detect cell transit events from transient increases in photon count rate (Fig. 2). The program reads all time tags from the Time Tagger device over USB 3.0 every 10 ms and generates decay histograms in between 1 ms clock pulses from the external function generator. The 1 ms period corresponds to 50,000 excitation pulses from the 50 MHz laser source. Each histogram records the distribution of time differences (dt) between PMT photon detection time tags and their preceding excitation laser sync pulse (Fig. 2A). The 1 ms histogramming period is small enough that each cell transit event (with a nominal 5 ms transit duration) results in about 5 consecutive histograms with significant autofluorescence photon counts. As such, the program constantly compares the photon counts of the 1 ms histograms against the background photon count rate. The background photon count rate threshold is estimated as the median value of the 1ms histogram photon counts over the last 5 seconds, given the low duty cycle of a cell transit event (typically 1% to 10% depending on sample concentration) over any given 5 seconds. A logical binary stream of cell detection decisions is generated by this threshold over background (Fig. 2B). If more than two consecutive histograms have higher than a user-defined multiple (typically 1.5x) of this threshold for background photon counts, a cell transit event is identified, and these histograms are combined to produce and record the associated autofluorescence decay histogram.

**Fig. 2:**
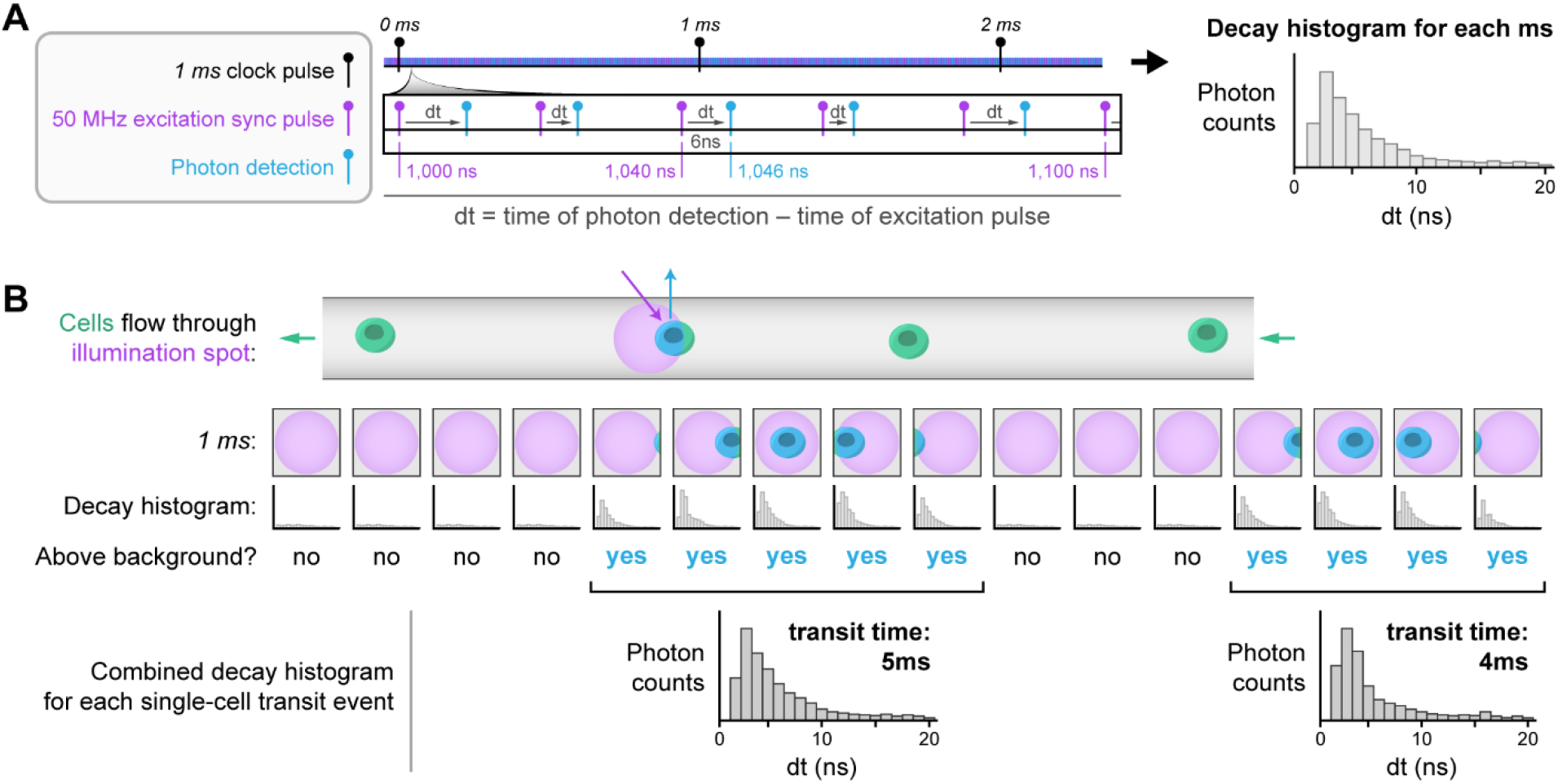
Custom photon stream TCSPC histogramming and cell transit event detection algorithm. (A) The time tagger device records dt, which is the photon detection time relative to the excitation laser sync pulse. Multiple dt values are recorded as a histogram in between clock pulses from a 1 ms external clock. The slow 1 ms clock keeps track of macro-time events such as transit of a single cell through the laser illumination spot. (B) The photon counts within the stream of 1 ms histograms are compared against the background count rate to produce a logical binary stream of cell detection decisions. If more than two consecutive 1 ms histograms have higher than background counts, these histograms are combined to produce the autofluorescence decay histogram assigned to a single cell transit event.

### 2.4 Real-time phasor analysis

Phasor representation [31] is a computationally low-cost and fast way to reduce each single cell decay histogram to a single point on a two-dimensional phasor plot that provides contrast between different fluorescence lifetimes. Briefly, the phasors are the complex coefficients of the Fourier series expansion of the time-resolved decay at harmonics of the laser repetition frequency. Typically, the first harmonic carries the most information about the decay and subsequent harmonics provide progressively less information about the higher-order oscillations in the decay. Therefore, the first harmonic phasor was calculated and plotted in real time for each cell as a (G, S) Cartesian coordinate pair according to 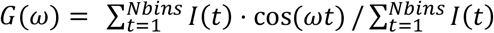 and 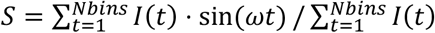, where ω = 2*πnf* is the frequency of phasor calculation with *f* being the laser repetition frequency (50 MHz here) and *n* being the harmonic number (one here).

Phasor analysis has the linearity property, such that a combination of two fluorescent molecular species gives rise to a linear combination in the phasor space, with the combined phasor falling on the straight line connecting the phasors of the individual constituting species [32].

Deconvolution of the impulse (or the instrument) response function (IRF) in the phasor space amounts to a scale and rotation transformation of the raw phasor coordinates [33]. The magnitude and phase of this transformation was estimated from the raw phasor of the time-resolved IRF curve, which was measured from a specular reflection of the excitation laser spot by a retroreflector mirror, and had a 350 ps FWHM.

After plotting the single-cell phasor points for an initial flow run, the program asks the user to draw a polygon gate over the phasor plot to select a subpopulation of interest. In subsequent flow runs, the program generates a real-time sort decision based on whether new cells fall within this phasor gate. While we have not implemented a physical sorting mechanism for our flow cell in this study, the ability to control such a mechanism in real time using the phasor gating sort decision is implemented.

### 2.5 Post-processing lifetime fit analysis

The recorded time-resolved single-cell decay histograms were post processed in Matlab 2021b (MathWorks, Natick, MA) using an iterative re-convolution algorithm that fits the measured decays to a bi-exponential model *I*(*t*) = *I*_0_ ⋅ *α*_1_ exp (*t*/*τ*_1_) *α*_2_ exp (*t*/*τ*_2_) *offset* ∗ *IRF*(*t*), where ∗ denotes convolution, by minimizing the sum of squared residuals over all time bins. A summary parameter known as the mean fluorescence lifetime is calculated as *τ*_m_ = *α*_1_*τ*_1_ *α*_2_*τ*_2_. Despite the higher computational cost, these fit parameters (*α*_1_, *τ*_1_, *α*_2_, *τ*_2_, *τ*_m_) typically capture most of the relevant information contained in the time-resolved autofluorescence decay curves including the lifetime of free (*τ*_1_) and protein-bound (*τ*_2_) conformations of NAD(P)H, and the fractional contribution of either conformation (*α*_1_, *α*_2_) to the total decay.

Where applicable, a uniform manifold approximation and projection (UMAP [34]) was employed in Python (Scikit-learn [35]) for dimension reduction of the intensity and lifetime parameters to produce 2D scatter plots that maintain the distance between cells in the original multidimensional feature space. To quantify the utility of these phasor- or fit-derived lifetime parameters for discriminating the metabolic phenotype of the cells, random forest classifiers where trained and tested (with a 50:50 train to test data split) for their ability to correctly identify treatment or culture conditions. Receiver operating characteristic (ROC) curves were generated and the area under the curve (AUC) was reported.

### 2.6 Samples and cells for flow experiments

Fluorescent polystyrene microspheres (beads) were used to validate the accuracy of lifetime measurements with the flow system. Two types of blue fluorescent beads, 10 µm Bright Blue (ex 360 nm, em 407 nm, Polysciences Inc., Warrington, PA) and 5 µm Glacial Blue (ex 360 nm, em 420 nm, Bangs Laboratories Inc., Fishers, IN) were used to prepare diluted stock suspensions of roughly 1 million beads per milliliter in deionized (DI) water according to the manufacturer labels. Five mixtures of the two stocks (with volumetric ratios of 1:5, 1:2, 1:1, 2:1, 5:1) were prepared for flow experiments. The true bead mixture ratios and the fluorescence lifetimes of the two beads were determined using two-photon (2P) excited FLIM at 750 nm excitation, 440/80 nm emission. 2P FLIM was collected with an Ultima multiphoton microscope (Bruker Nano Inc., Madison, WI) equipped with SPC-150 timing electronics (Becker & Hickl GmbH, Berlin, Germany) [26] and fit to a single-exponential decay in SPCImage 8.8 (Becker & Hickl GmbH, Berlin, Germany). A 40× water immersion 1.13 NA objective lens (Nikon) and 60 seconds integration time per field of view was used.

Jurkat T cells, an immortalized line of human T lymphocyte cells from a leukemia patient, were cultured in Advanced RPMI 1640 (Gibco) + 10% fetal bovine serum (Sigma Aldrich) + 1% penicillin/streptomycin (Gibco) and maintained at 5% CO_2_ and 37°C prior to flow runs. These cells were used for testing the sensitivity of the lifetime flow system to the effects of a known oxidative metabolism inhibitor, sodium cyanide. Briefly, 4 mM solution of sodium cyanide (Sigma-Aldrich) in DPBS (Gibco) was prepared from powder. For treatment, cells were centrifuged at 1200 rpm for 5 minutes to form a pellet. The media was aspirated, and the pellet was washed once with DPBS before being resuspended in fresh DPBS for the control condition, or in DPBS plus 4 mM sodium cyanide for the treatment condition, at a concentration of 100,000 cells per milliliter, 15 minutes prior to flow experiments.

Primary human T cells were isolated from the peripheral blood of a healthy donor using a negative selection kit (EasySep, STEMCELL Technologies Canada Inc., Vancouver, BC) and used for testing the sensitivity of the lifetime flow system to T cell activation. Informed consent was obtained in accordance with guidelines set forth by the Institutional Review Board of the University of Wisconsin-Madison. The T cells were cultured for 48 hours in ImmunoCult medium (STEMCELL Technologies Canada Inc., Vancouver, BC) for the control condition, or in ImmunoCult medium + 25µL/mL CD3/CD28/CD2 T cell activator (STEMCELL) for the activated condition. The cells were resuspended in DPBS as before, at a concentration of 1 million/mL, 15 minutes prior to flow experiments.

Primary neural stem cells (NSC) were isolated from the hippocampus of adult C57BL/6J (Jackson Laboratory) mouse brains as described previously [36]. All facilities used for maintaining the colony of mice used in this study have been approved by the Research Animal Resources and Compliance (RARC) at UW-Madison. Activated NSCs were cultured as previously described at 37°C/5% CO_2_ in serum-free media (aNSC media): DMEM/F12 GlutaMax (Invitrogen 10565018) with B27 (1:50, Invitrogen 17504044), penicillin-streptomycin-fungizone (1:100, Invitrogen 15140122), and 20 ng/mL FGF-2 and EGF (PeproTech 100-18B and AF-100-15). To generate quiescent NSCs, aNSCs were plated onto PLO- and laminin-coated plates and treated with BMP-4, with FGF-2 and without EGF (qNSC media), using previously described protocols [37–41]. qNSC media is similar to aNSC media except for the removal of EGF and the addition of 50 ng/mL BMP-4 (Fisher Scientific 5020BP010). For these experiments, both aNSC and qNSCs were plated into either PLO-laminin-coated ibidi 8 well chamber, or 6 well plates (ibidi GmbH, Gräfelfing, Germany). After initial induction of quiescence, qNSCs were fed at least once every two days and were considered quiescent after 4 days of qNSC media treatment. aNSCs and qNSCs in 6 well plates were trypsinized using the following protocol: Media was removed, and wells were treated with 0.05% trypsin (Invitrogen 25300-054) at 37°C for 5 minutes. Cells were then quenched with media, mechanically triturated, and pelleted in a centrifuge by spinning at 120xg for 4 minutes. Single cells were then suspended in DPBS as before, at a concentration of 2 million/mL, 15 minutes prior to flow experiments. The same day, 2P FLIM of the aNSCs and qNSCs cultured in ibidi 8 well chambers was performed with the Ultima multiphoton microscope (Bruker Nano Inc., Madison, WI) equipped with SPC-150 timing electronics (Becker & Hickl GmbH, Berlin, Germany) [26] and fit to a two-exponential decay in SPCImage 8.8 (Becker & Hickl GmbH, Berlin, Germany). A 40× water immersion 1.13 NA objective lens (Nikon) and 60 seconds integration time per field of view was used. NAD(P)H lifetime was excited at 750nm and collected at 440/80nm, while punctate autofluorescence (PAF) intensity, detected in a subpopulation of lysosomes, was collected at 590/50nm. These signals have been previously shown to separate qNSCs and aNSCs [36].

### 2.7 Statistical analysis

Statistical analysis and generation of scatter plots was done in Matlab. Unpaired (*i.e*., two-sample) *t*-test was used to determine if there is a statistically significant difference between population means and the *p*-value range was reported (****, *p* < 0.0001). Violin plots [42] were superimposed with boxplots showing the interquartile range with whiskers extending from the box to 1.5 times the interquartile range. White dots show the median and horizontal lines show the mean.

#### 3. Results

### 3.1 Discrimination of beads based on fluorescence lifetime

Fluorescent beads were first used as a known standard to test the accuracy of the fluorescence lifetime flow cytometer. The ground truth for fluorescence lifetime of the blue beads and their true mixture ratio was established by 2P FLIM as shown in Fig. 3A. The BB beads have a lifetime of τ = 2084 ps and the GB beads have a lifetime of τ = 1684 ps. The five bead mixtures were flowed through the fluorescence lifetime flow cytometer over 300 seconds of flow time each. Time-resolved fluorescence decays from an average of 1500 beads per flow run were recorded. A polygon phasor gate was drawn on the population of beads with the shorter lifetime to identify them as GB while beads falling outside the gate were identified as BB (Fig. 3B). The post-processing fit analysis of the time-resolved decays measured by the flow system determined a lifetime of τ = 2100 ps for BB beads and a lifetime of τ = 1720 ps for GB beads, which is in good agreement with the 2P FLIM values. The ratio of beads counted in this fashion was compared against the true mixture ratio for each preparation. The resulting regression line presented in Fig. 3C shows good agreement and a linear relationship with a coefficient of determination of 0.999 and a slope of 1.07, which is close to one.

**Fig. 3:**
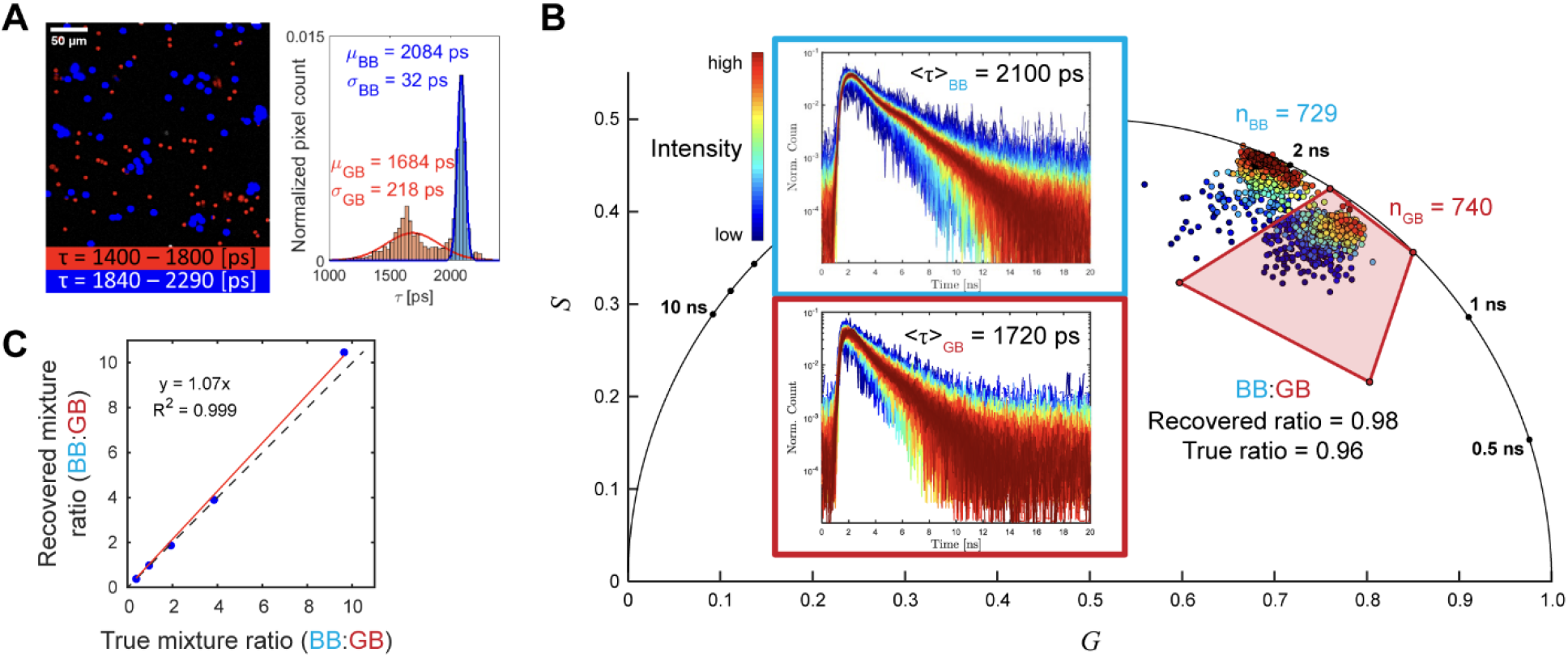
The fluorescence lifetime flow cytometer can discriminate blue fluorescent beads based on their lifetimes. (A) Two types of blue fluorescent beads, 10 µm Bright Blue (BB, τ = 2084ps) and 5 µm Glacial Blue (GB, τ = 1684ps) were imaged using two-photon (2P) FLIM and their single-exponential fluorescence lifetimes were determined. (B) Suspensions of the two beads with different mixture ratios were flowed through the fluorescence lifetime flow cytometer; the bead types were identified and counted based on gating of the phasor representation of their measured fluorescence decays. On average, 1500 beads were counted in each of the five 5-minute flow runs. Insets show the time-resolved decays measured from the beads color-coded by the photon count (*i.e*., intensity) of each bead. The fluorescence lifetimes of the beads measured by the fluorescence lifetime flow cytometer (BB, τ = 2100ps; GB, τ = 1720ps) agree with the standard 2P FLIM measurements. (C) The recovered mixture ratio estimated by the fluorescence lifetime flow cytometer closely matches the true mixture ratio of the preparations in five experiments, forming a regression line with a slope of 1.07 and a coefficient of determination of 0.999.

### 3.2 Sensitivity to metabolic inhibition in Jurkat T cells

Cyanide is an inhibitor of the electron transport chain, and the resulting imbalance towards glycolysis causes an increase in the free NAD(P)H pool. This experiment tested the sensitivity of the fluorescence lifetime flow cytometer to this known metabolic shift. Jurkat T cells treated with 4 mM sodium cyanide for 15 minutes present with shorter NAD(P)H autofluorescence lifetime (phasor position shifted to lower right) than control cells as shown in the phasor plot in Fig. 4A. Post-processing fit analysis of the time-resolved decays shows cyanide-treated Jurkat T cells have shorter NAD(P)H mean lifetime (τ_m_) (Fig. 4B), higher fraction of free NAD(P)H (α_1_) (Fig. 4C), and shorter free NAD(P)H lifetime (τ_1_) (Fig. 4D) than control cells, while bound NAD(P)H lifetime (τ_2_) is not significantly different (Fig. 4E). These results are consistent with prior 2P FLIM measurements of NAD(P)H lifetimes in cyanide-treated cells, and with the expected effects of cyanide inhibition of the mitochondrial electron transport chain [6,8,43,44].

**Fig. 4:**
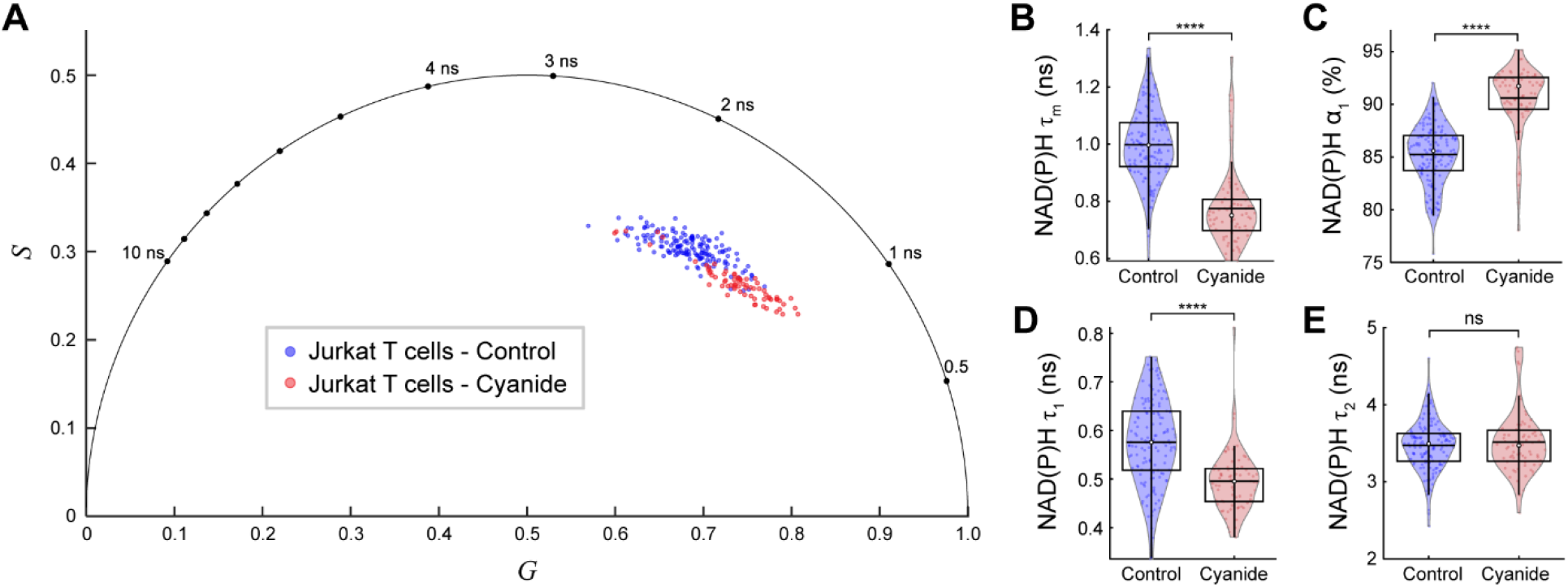
The fluorescence lifetime flow cytometer is sensitive to metabolic perturbation in Jurkat T cells. (A) Phasor representation of the Jurkat T cells NADH(P)H lifetime measured in flow. The control T cells were resuspended in DPBS flow medium, and the treated T cells were resuspended in 4mM sodium cyanide in DPBS, 15 minutes prior to the flow experiment. The cyanide-treated cells cluster separately from the control cells and present with a shorter fluorescence lifetime, as expected [8]. Post-processing analysis of the time-resolved decay was also performed by fitting to a bi-exponential decay model. Cyanide-treated cells have (B) shorter NAD(P)H mean fluorescence lifetime (τ_m_), (C) increased free fraction of NAD(P)H (α_1_), and (D) shorter free NAD(P)H lifetime (τ_1_), while (E) the bound NAD(P)H lifetime (τ_2_) is not significantly different. Cyanide-treated T cells, n = 71. Control T cells, n = 140. ****, unpaired *t*-test *p* < 0.0001. Box plots show the interquartile range and whiskers extend from the box to 1.5 times the interquartile range. White dots show the median and red horizontal lines show the mean.

### 3.3 Sensitivity to activation of primary human T cells

To demonstrate the application of the fluorescence lifetime flow cytometer to T cell manufacturing, activated and quiescent primary human T cells were assessed, as T cell activation is known to increase glycolysis and change the NAD(P)H lifetime [45–47]. The NAD(P)H phasor plot from the fluorescence lifetime flow cytometer (Fig. 5A) shows that activated T cells cluster separately, with a shorter fluorescence lifetime, from quiescent T cells. Residual red blood cells in the sample present with the very short (τ < 100 ps) fluorescence lifetime of hemoglobin photoproducts [48] and, depending on their axial position during transit through the illumination spot (in or out of focus) and contribution of background signal, appear on the axis connecting the zero lifetime (*i.e*., IRF) position and the phasor location of the background fluorescence (measured from the DPBS resuspension medium in the channel). Post-processing fit analysis of the time-resolved T cell decays measured in flow (Fig. 5B-E) shows shorter NAD(P)H τ_m_, τ_1_, and τ_2_, and higher NAD(P)H α_1_ for activated T cells compared to quiescent T cells. UMAP projection of the NAD(P)H lifetime fit parameters (τ_m_, τ_1_, τ_2_, α_1_) and phasors (1^st^, 2^nd^ harmonics) shows separate clustering of activated and quiescent T cells (Fig. 5F). Random forest classifiers trained on either NAD(P)H phasors or lifetime fit parameters can correctly identify the activation culture condition of the T cells with high accuracy as shown by the high area under the curve (AUC) for their receiver operating characteristic (ROC) curves (Fig. 5G). Trypan blue exclusion showed viability of 84% in flowed cells with exposure to the laser beam and 89% in cells that were not flowed.

**Fig. 5:**
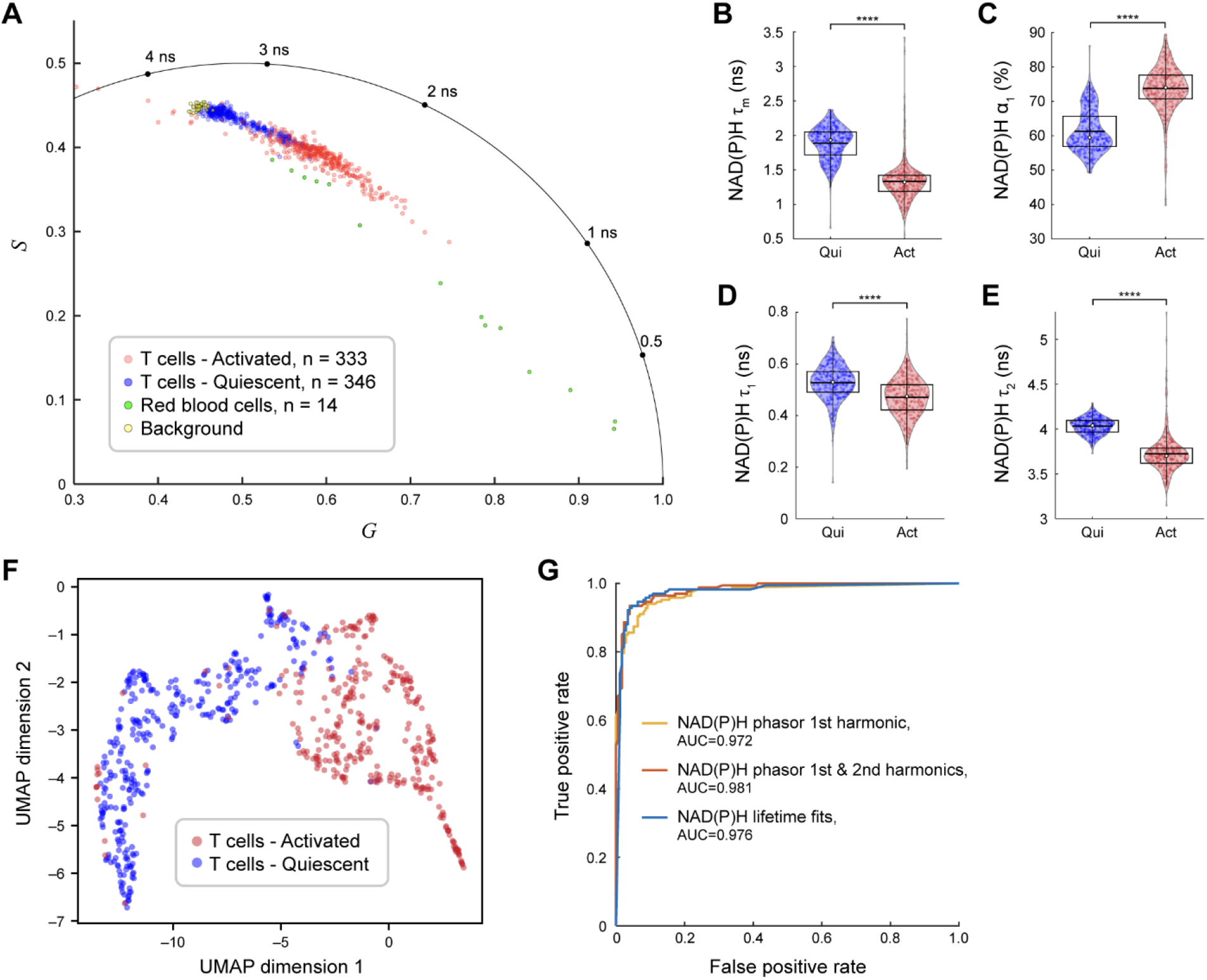
The fluorescence lifetime flow cytometer is sensitive to activation state of primary T cells. Primary T cells were isolated from peripheral blood of a healthy donor and cultured for 48 hours in Immunocult medium for the control quiescent condition, or Immunocult medium + tetrameric antibody against CD3/CD28/CD2 for the activated condition. Both quiescent and activated T cells were resuspended in DPBS flow medium 15 minutes prior to the flow experiments. (A) Phasor representation of NAD(P)H lifetime from the primary human T cells measured in flow. The activated cells cluster separately from the control quiescent cells and present with a shorter fluorescence lifetime. Red blood cells have shorter fluorescence lifetimes, and DPBS background has a longer fluorescence lifetime. Post-processing analysis of the time-resolved decays were also performed by fitting to a bi-exponential decay model. Activated cells have (B) shorter NAD(P)H τ_m_, (C) increased NAD(P)H α_1_, (D) shorter NAD(P)H τ_1_, and (E) shorter NAD(P)H τ_2_. (F) UMAP projection of the NAD(P)H lifetime fit parameters (τ_m_, τ_1_, τ_2_, α_1_) and phasors (1^st^, 2^nd^ harmonics) shows separate clustering of the quiescent and activated T cells. (G) Random forest classifiers trained on NAD(P)H phasor or lifetime parameters (50% training; 50% testing split) can predict the activation state of the T cells with good accuracy as demonstrated by the area under the curve (AUC) of the receiver operating characteristic (ROC) curves. Activated T cells, n = 333. Quiescent T cells, n = 346. ****, unpaired *t*-test *p* < 0.0001. Box plots show the interquartile range and whiskers extend from the box to 1.5 times the interquartile range. White dots show the median and red horizontal lines show the mean.

### 3.4 Sensitivity to activation of primary neural stem cells

To benchmark the measurements of our flow system against a standard technique, aNSCs and qNSCs were imaged on the two-photon FLIM microscope over 2 hours, resulting in an image dataset containing 1551 cells (n=651 aNSCs and n=900 qNSCs). Post-processing analysis of these images (including single cell segmentation, pixel-wise lifetime fitting, and summarizing pixel value averages over cell masks) required an additional day of analysis time. Representative NAD(P)H τ_m_ and PAF intensity images are shown in Fig. 6A,B. aNSCs present with shorter NAD(P)H τ_m_, τ_1_, and τ_2_, and higher NAD(P)H α_1_ than qNSCs (Fig.6C-F), while qNSCs have higher PAF intensity (Fig. 6G). UMAP projection of NAD(P)H lifetime and PAF intensity parameters shows clear separation of aNSCs and qNSCs (Fig. 6H). The ROC curves (Fig. 6I) for random forest classifiers trained on PAF intensity, NAD(P)H lifetime fit parameters, or both show that intensity values alone do not provide good classification accuracy (AUC = 0.779), while NAD(P)H lifetime parameters provide good classification accuracy (AUC = 0.985) and both PAF intensity and NAD(P)H lifetime parameters provide near-perfect classification accuracy (AUC = 0.996).

**Fig. 6:**
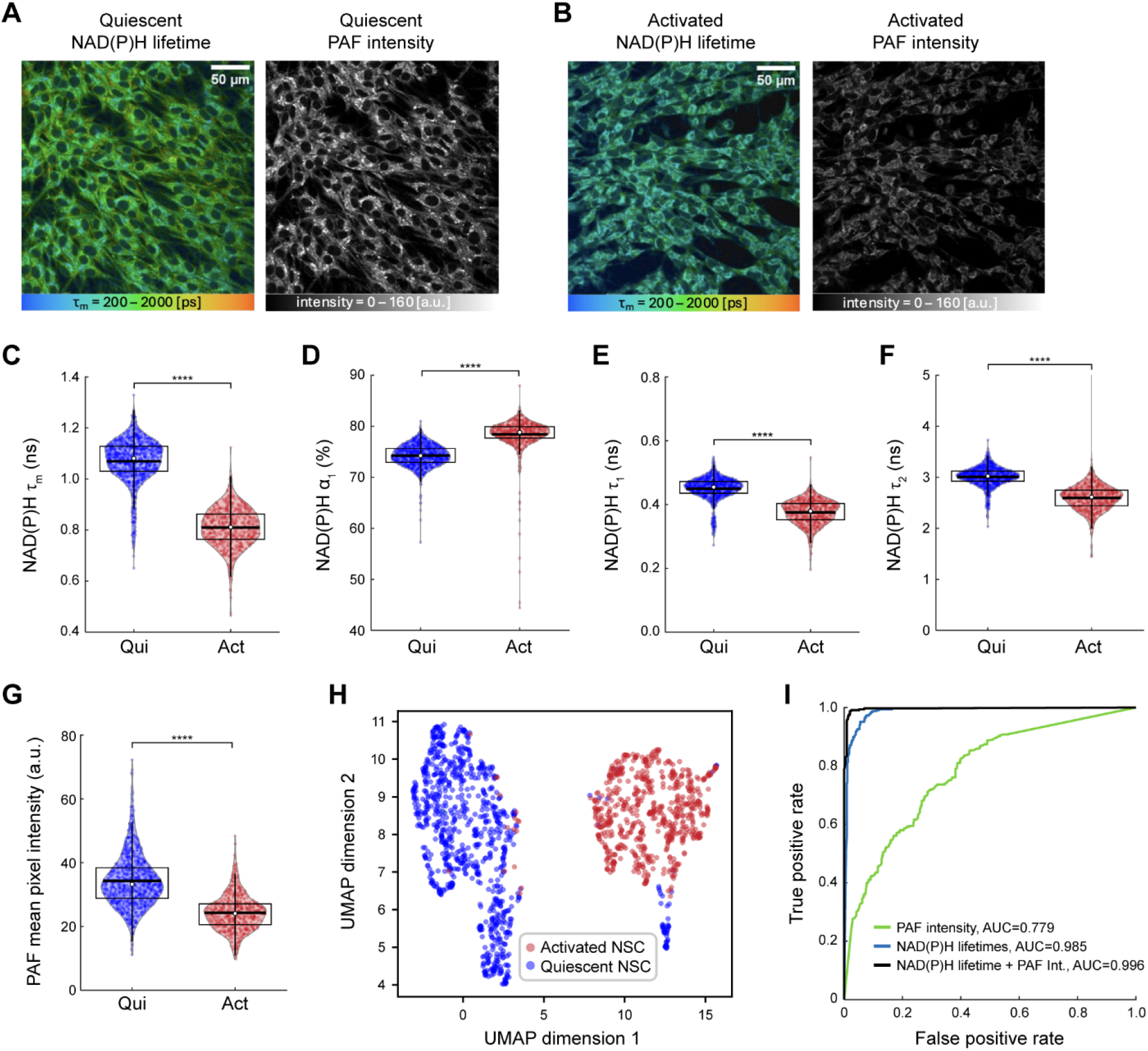
Two-photon FLIM is sensitive to the activation state of NSCs. Representative 2P-excited NAD(P)H lifetime and punctate autofluorescence (PAF = 590/50nm) intensity images of primary mouse NSCs in (A) quiescent and (B) activating culture conditions. The qNSCs present with longer NAD(P)H lifetime and higher PAF intensity compared to aNSCs. Fitting analysis of the NAD(P)H lifetime images show that aNSCs present with (C) shorter NAD(P)H τ_m_, (D) higher NAD(P)H α_1_, (E) shorter NAD(P)H τ_1_, and (F) shorter NAD(P)H τ_2_. (G) PAF intensity is higher in qNSCs. (H) UMAP projection of the NAD(P)H lifetime fit parameters (τ_m_, τ_1_, τ_2_, α_1_) and PAF intensity shows separate clustering of qNSCs and aNSCs. (I) Random forest classifiers trained on PAF intensity and NAD(P)H lifetime parameters (50% training; 50% testing split) can predict the activation state of the NSCs with good accuracy as demonstrated by the AUC of ROC curves. qNSCs, n = 900. aNSCs, n = 651. ****, unpaired *t*-test *p* < 0.0001. Box plots show the interquartile range and whiskers extend from the box to 1.5 times the interquartile range. White dots show the median and red horizontal lines show the mean.

aNSCs and qNSCs from the same culture were resuspended in DPBS and flowed through the fluorescence lifetime flow cytometer to acquire autofluorescence with the 375nm pulsed laser in two spectral channels: 440/80nm for NAD(P)H lifetime and 590/50nm for PAF intensity. Two 5-minute flow runs for the aNSCs and qNSCs resulted in acquisition of 1497 cells (n=920 aNSCs and n=577 qNSCs). The NAD(P)H phasor plot (Fig. 7A) shows separation of aNSCs with a shorter fluorescence lifetime than qNSCs. The same NAD(P)H phasor plot overlaid with a colormap corresponding to the intensity of PAF autofluorescence in the second spectral channel shows a positive correlation between NAD(P)H fluorescence lifetime and PAF intensity (Fig. 7B). Post-processing fit analysis of the time-resolved NSC decays acquired in flow shows the same trends in lifetime fit parameters that was observed with 2P FLIM. That is, aNSCs present with shorter NAD(P)H τ_m_, τ_1_, and τ_2_, and higher NAD(P)H α_1_ than qNSCs (Fig.7C-F), while qNSCs have higher PAF intensity (Fig. 7G). UMAP projection of PAF intensity, NAD(P)H phasor, and NAD(P)H lifetime fit parameters shows clear separation of aNSCs and qNSCs (Fig. 7H). The ROC curves (Fig. 7I) for random forest classifiers trained on PAF intensity, NAD(P)H phasor, or both PAF intensity and NAD(P)H lifetime fit parameters show that intensity values alone do not provide good classification accuracy (AUC = 0.708), while NAD(P)H phasor provides good classification accuracy (AUC = 0.935) and both PAF intensity and NAD(P)H lifetime parameters provide excellent classification accuracy (AUC = 0.959).

**Fig. 7:**
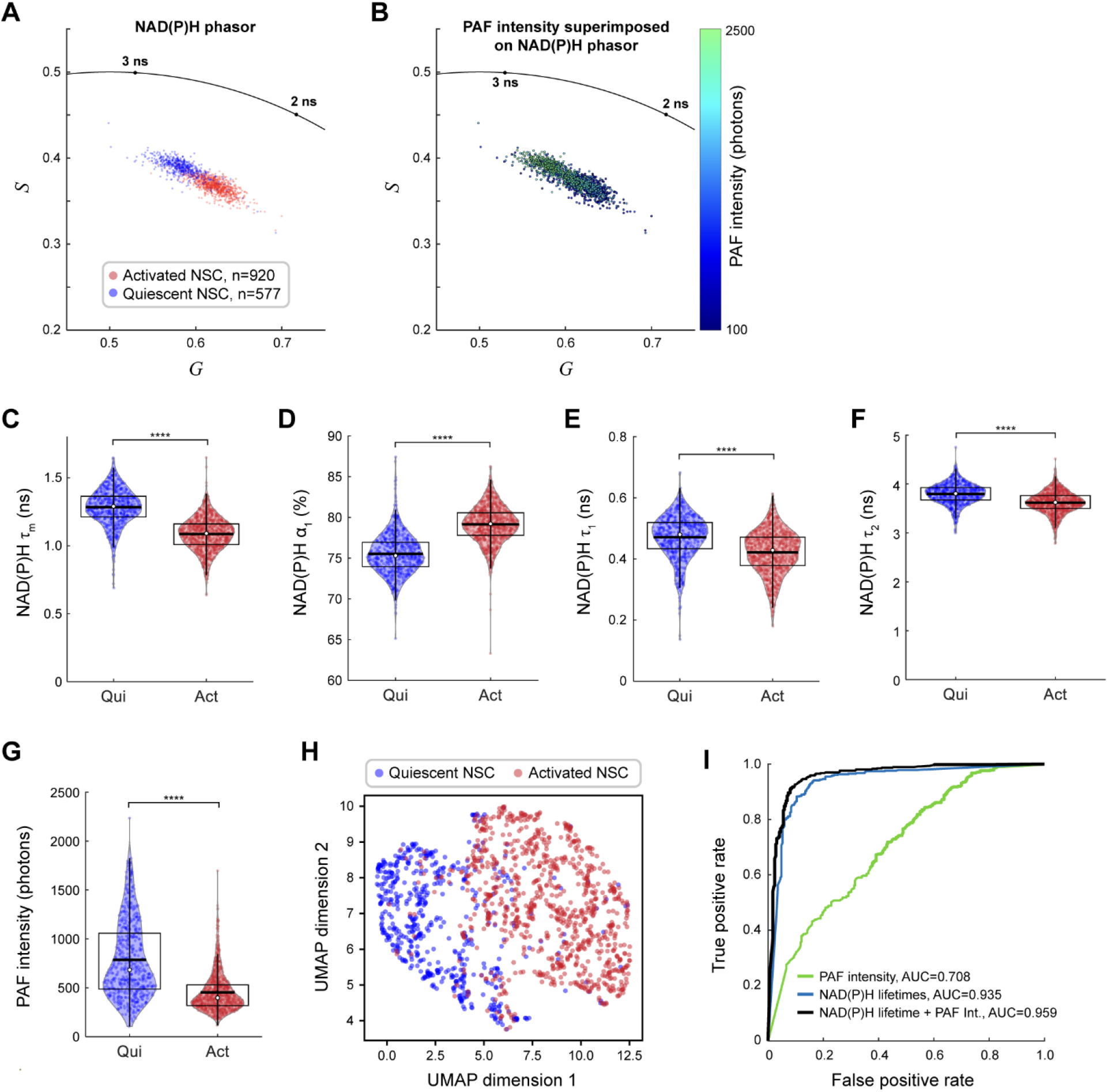
The fluorescence lifetime flow cytometer is sensitive to the activation state of NSCs. (A) Phasor representation of the NAD(P)H decays measured in flow (by PMT 1) from primary NSCs in quiescent and activating culture conditions. The qNSCs present with longer NAD(P)H lifetime and cluster separately from aNSCs. (B) The same phasor plot with a superimposed colormap corresponding to the intensity of the PAF channel (590/50 nm) shows a positive correlation between long NAD(P)H lifetime and high PAF intensity. In agreement with 2P FLIM, fitting analysis of the decays acquired by the fluorescence lifetime flow cytometer shows aNSCs present with (C) shorter NAD(P)H τ_m_, (D) higher NAD(P)H α_1_, (E) shorter NAD(P)H τ_1_, and (F) shorter NAD(P)H τ_2_. (G) The PAF intensity is higher in qNSCs. (H) UMAP projection of the NAD(P)H phasor coordinates, NAD(P)H lifetime fit parameters (τ_m_, τ_1_, τ_2_, α_1_), and PAF intensity (all measured by the flow system) shows separate clustering of qNSCs and aNSCs. (I) Random forest classifiers trained on PAF intensity and NAD(P)H lifetime parameters (50% training; 50% testing split) can predict the activation state of the NSCs with good accuracy as demonstrated by AUC of the ROC curves. qNSCs, n = 577. aNSCs, n = 920. ****, unpaired *t*-test *p* < 0.0001. Box plots show the interquartile range and whiskers extend from the box to 1.5 times the interquartile range. White dots show the median and red horizontal lines show the mean.

## 4. Discussion

We presented a single photon-excited autofluorescence lifetime flow cytometer using the sensitive time-correlated single photon counting (TCSPC) technique for time-resolved acquisition of autofluorescence decays from single cells in a microfluidic flow cell. This system allows for label-free, non-destructive, longitudinally repeatable measurements with real-time analysis in a flow geometry that is amenable to preclinical, clinical, and cell manufacturing settings that require automation, higher-throughput, and unaltered cell state.

The acquisition speed of this system is inversely proportional to the target number of photons per cell. At our target for accurate two-component lifetime analysis (10,000 photons or 5 ms interrogation per cell [27,28]), the upper limit of acquisition speed is 100 cells per second, assuming a 50% cell transit duty cycle. This throughput is one or two orders of magnitude lower than conventional flow cytometry. However, this trade-off in speed is justified by the advantages of label-free metabolic analysis, which removes the need for reagents, time, and labor associated with labeling cells, and avoids any unwanted influence of exogenous reagents on cell function or clinical release [49–51].

When compared to existing alternatives that perform time-resolved autofluorescence measurements, *i.e*., confocal, or multiphoton laser-scanning FLIM, our flow system provides nearly 100x throughput advantage, real-time phasor analysis, and a simplified post-processing analysis pipeline. As a specific example, we compare the acquisition and analysis of the data in Figs. 6 and 7 using a 2P FLIM microscope and our fluorescence lifetime flow system, respectively. Two hours of active imaging by an expert user on the multiphoton microscope resulted in images of 1551 NSCs (at near 100% confluency) which translates to an acquisition time of 4.6 seconds per cell. Over the next day, the FLIM images were analyzed by performing pixel-wise decay fitting using one software tool (SPCImage 8.8, Becker & Hickl GmbH), followed by single-cell segmentation using another software tool (Cellpose [52]), followed by averaging of pixel lifetimes over cell masks using a custom script (Cell Analysis Tools [53]). All steps required hands-on user input. In contrast, once the cells were loaded into a syringe, the flow system acquired single-cell decays from 1497 NSCs in 10 minutes, without any user input. This translates to an acquisition time of 0.4 seconds per cell, which is 12x faster than the 2P FLIM. The cell transit duty cycle in this experiment was 6.25%, which means the throughput could have been even higher with a more concentrated sample (*e.g*., 96x faster than 2P FLIM with 50% cell transit duty cycle). The phasor plots were generated by the system in real time and saved along with the raw single-cell decays. Post-processing analysis was performed by running a single Matlab lifetime fitting script pointed to the raw data folder, requiring no user input. Since only whole-cell decays were saved, no image segmentation or pixel averaging step was needed. The entire post-processing analysis of the flow data required about 10 minutes.

From an instrumentation perspective, this fluorescence lifetime flow cytometer has simplified optics, a smaller footprint, and uses lower-cost single-photon excitation lasers compared to multiphoton FLIM, while maintaining high accuracy in fluorescence decay measurements and high sensitivity to cell function. The picosecond-pulsed UV diode laser in our system costs about $15,000 which is over 10x lower than a femtosecond-pulsed Ti:Sapphire laser, power supply, water chiller, polarization optics, and Pockels cell needed for a 2P FLIM system. The flow geometry removes the need for scanning optics (scan lens, galvanometer mirrors, and electronics). As such, the flow system has an area footprint over 5x smaller than a 2P FLIM system.

While we limited our analysis to binary classification based on culture condition, the spread of NAD(P)H lifetime or phasor parameters shows that metabolic inhibition, T cell activation, and stem cell activation span a continuous spectrum. A continuous readout of cell metabolic state based on autofluorescence lifetimes may provide more insight into cell function than a binary readout based on surface markers (*e.g*., CD69), particularly when assessing cell fitness for cell therapy. The flow geometry and real-time phasor analysis and sort decision in this system will enable autofluorescence lifetime-based cell sorting or enrichment in the future, which will further broaden the applicability of this real-time single-cell metabolic assessment technique.

## Acknowledgements

We thank the Fab Lab at the Morgridge Institute for Research for technical assistance, and Matthew Stefely for figure illustrations.

## References

1. Chance B, Schoener B, Oshino R, Itshak F, Nakase Y. Oxidation-reduction ratio studies of mitochondria in freeze-trapped samples. NADH and flavoprotein fluorescence signals. Journal of Biological Chemistry 1979;254:4764–4771.

2. Lakowicz JR, Szmacinski H, Nowaczyk K, Johnson ML. Fluorescence lifetime imaging of free and protein-bound NADH. Proceedings of the National Academy of Sciences 1992;89:1271–1275.

3. Kalinina S, Breymayer J, Schäfer P, Calzia E, Shcheslavskiy V, Becker W, Rück A. Correlative NAD(P)H-FLIM and oxygen sensing-PLIM for metabolic mapping. Journal of Biophotonics 2016;9:800–811.

4. Meleshina AV, Dudenkova VV, Bystrova AS, Kuznetsova DS, Shirmanova MV, Zagaynova EV. Two-photon FLIM of NAD(P)H and FAD in mesenchymal stem cells undergoing either osteogenic or chondrogenic differentiation. Stem Cell Res Ther 2017;8:15.

5. Rice WL, Kaplan DL, Georgakoudi I. Two-Photon Microscopy for Non-Invasive, Quantitative Monitoring of Stem Cell Differentiation. PLOS ONE 2010;5:e10075.

6. Huang S, Heikal AA, Webb WW. Two-Photon Fluorescence Spectroscopy and Microscopy of NAD(P)H and Flavoprotein. Biophysical Journal 2002;82:2811–2825.

7. Scott TG, Spencer RD, Leonard NJ, Weber G. Emission properties of NADH. Studies of fluorescence lifetimes and quantum efficiencies of NADH, AcPyADH, and simplified synthetic models. J. Am. Chem. Soc. 1970;92:687–695.

8. Bird DK, Yan L, Vrotsos KM, Eliceiri KW, Vaughan EM, Keely PJ, White JG, Ramanujam N. Metabolic Mapping of MCF10A Human Breast Cells via Multiphoton Fluorescence Lifetime Imaging of the Coenzyme NADH. Cancer Research 2005;65:8766–8773.

9. Skala MC, Riching KM, Gendron-Fitzpatrick A, Eickhoff J, Eliceiri KW, White JG, Ramanujam N. In vivo multiphoton microscopy of NADH and FAD redox states, fluorescence lifetimes, and cellular morphology in precancerous epithelia. Proceedings of the National Academy of Sciences 2007;104:19494–19499.

10. Sharick JT, Favreau PF, Gillette AA, Sdao SM, Merrins MJ, Skala MC. Protein-bound NAD(P)H Lifetime is Sensitive to Multiple Fates of Glucose Carbon. Sci Rep 2018;8:5456.

11. Pires L, Nogueira MS, Pratavieira S, Moriyama LT, Kurachi C. Time-resolved fluorescence lifetime for cutaneous melanoma detection. Biomed. Opt. Express, BOE 2014;5:3080–3089.

12. Kapsokalyvas D, Barygina V, Cicchi R, Fiorillo C, Pavone FS. Evaluation of the oxidative stress of psoriatic fibroblasts based on spectral two-photon fluorescence lifetime imaging. In: Multiphoton Microscopy in the Biomedical Sciences XIII. Vol 8588. SPIE; 2013. p 386–395.

13. Nickel AG, von Hardenberg A, Hohl M, Löffler JR, Kohlhaas M, Becker J, Reil J-C, Kazakov A, Bonnekoh J, Stadelmaier M, Puhl S-L, Wagner M, Bogeski I, Cortassa S, Kappl R, Pasieka B, Lafontaine M, Lancaster CRD, Blacker TS, Hall AR, Duchen MR, Kästner L, Lipp P, Zeller T, Müller C, Knopp A, Laufs U, Böhm M, Hoth M, Maack C. Reversal of Mitochondrial Transhydrogenase Causes Oxidative Stress in Heart Failure. Cell Metabolism 2015;22:472–484.

14. Qian T, Heaster TM, Houghtaling AR, Sun K, Samimi K, Skala MC. Label-free imaging for quality control of cardiomyocyte differentiation. Nat Commun 2021;12:4580.

15. Evans ND, Gnudi L, Rolinski OJ, Birch DJS, Pickup JC. Glucose-dependent changes in NAD(P)H-related fluorescence lifetime of adipocytes and fibroblasts in vitro: Potential for non-invasive glucose sensing in diabetes mellitus. Journal of Photochemistry and Photobiology B: Biology 2005;80:122–129.

16. Quinn KP, Leal EC, Tellechea A, Kafanas A, Auster ME, Veves A, Georgakoudi I. Diabetic Wounds Exhibit Distinct Microstructural and Metabolic Heterogeneity through Label-Free Multiphoton Microscopy. Journal of Investigative Dermatology 2016;136:342–344.

17. Chacko JV, Eliceiri KW. Autofluorescence lifetime imaging of cellular metabolism: Sensitivity toward cell density, pH, intracellular, and intercellular heterogeneity. Cytometry Part A 2019;95:56–69.

18. Alturkistany F, Nichani K, Houston KD, Houston JP. Fluorescence lifetime shifts of NAD(P)H during apoptosis measured by time-resolved flow cytometry. Cytometry Part A 2019;95:70–79.

19. Cao R, Jenkins P, Peria W, Sands B, Naivar M, Brent R, Houston JP. Phasor plotting with frequency-domain flow cytometry. Opt. Express, OE 2016;24:14596–14607.

20. Valentino S, Ortega-Sandoval K, Lucero S, Houston KD, Houston JP. Correlating NAD(P)H lifetime shifts to treatment of breast cancer cells: a metabolic screening study with time-resolved flow cytometry. In: Imaging, Manipulation, and Analysis of Biomolecules, Cells, and Tissues XXI. Vol 12383. SPIE; 2023. p 19–24.

21. Buschke DG, Squirrell JM, Ansari H, Smith MA, Rueden CT, Williams JC, Lyons GE, Kamp TJ, Eliceiri KW, Ogle BM. Multiphoton Flow Cytometry to Assess Intrinsic and Extrinsic Fluorescence in Cellular Aggregates: Applications to Stem Cells. Microscopy and Microanalysis 2011;17:540–554.

22. Buschke D g., Squirrell J m., Vivekanandan A, Rueden C t., Eliceiri K w., Ogle B m. Noninvasive sorting of stem cell aggregates based on intrinsic markers. Cytometry Part A 2014;85:353–358.

23. Sands B, Jenkins P, Peria WJ, Naivar M, Houston JP, Brent R. Measuring and Sorting Cell Populations Expressing Isospectral Fluorescent Proteins with Different Fluorescence Lifetimes. PLOS ONE 2014;9:e109940.

24. Zipfel WR, Williams RM, Christie R, Nikitin AY, Hyman BT, Webb WW. Live tissue intrinsic emission microscopy using multiphoton-excited native fluorescence and second harmonic generation. Proceedings of the National Academy of Sciences 2003;100:7075–7080.

25. Xu C, Williams RM, Zipfel W, Webb WW. Multiphoton excitation cross-sections of molecular fluorophores. Bioimaging 1996;4:198–207.

26. Becker W. The bh TCSPC Handbook: 10th Edition. Becker & Hickl GmbH; 2023. 1048 p.

27. Nasser M, Meller A. Lifetime-based analysis of binary fluorophores mixtures in the low photon count limit. iScience 2022;25:103554.

28. Bouchet D, Krachmalnicoff V, Izeddin I. Cramér-Rao analysis of lifetime estimations in time-resolved fluorescence microscopy. Opt. Express 2019;27:21239.

29. Wagner M, Weber P, Bruns T, Strauss WSL, Wittig R, Schneckenburger H. Light Dose is a Limiting Factor to Maintain Cell Viability in Fluorescence Microscopy and Single Molecule Detection. International Journal of Molecular Sciences 2010;11:956–966.

30. Application Programmer’s Interface — Time Tagger User Manual 2.16.2.0 documentation. Available at: https://www.swabianinstruments.com/static/documentation/TimeTagger/api/index.html.

31. Digman MA, Caiolfa VR, Zamai M, Gratton E. The Phasor Approach to Fluorescence Lifetime Imaging Analysis. Biophysical Journal 2008;94:L14–L16.

32. Ranjit S, Malacrida L, Jameson DM, Gratton E. Fit-free analysis of fluorescence lifetime imaging data using the phasor approach. Nat Protoc 2018;13:1979–2004.

33. Martelo L, Fedorov A, Berberan-Santos MN. Fluorescence Phasor Plots Using Time Domain Data: Effect of the Instrument Response Function. J. Phys. Chem. B 2015;119:10267–10274.

34. McInnes L, Healy J, Melville J. UMAP: Uniform Manifold Approximation and Projection for Dimension Reduction. 2020. Available at: http://arxiv.org/abs/1802.03426.

35. Pedregosa F, Varoquaux G, Gramfort A, Michel V, Thirion B, Grisel O, Blondel M, Prettenhofer P, Weiss R, Dubourg V, Vanderplas J, Passos A, Cournapeau D, Brucher M, Perrot M, Duchesnay É. Scikitlearn: Machine Learning in Python. Journal of Machine Learning Research 2011;12:2825–2830.

36. Morrow CS, Tweed K, Farhadova S, Walsh AJ, Lear BP, Roopra A, Risgaard RD, Klosa PC, Arndt ZP, Peterson ER, Chi MM, Harris AG, Skala MC, Moore DL. Autofluorescence is a biomarker of neural stem cell activation state. Cell Stem Cell 2024;0. Available at: https://www.cell.com/cell-stem-cell/abstract/S1934-5909(24)00054-7.

37. Mira H, Andreu Z, Suh H, Lie DC, Jessberger S, Consiglio A, San Emeterio J, Hortigüela R, Marqués-Torrejón MÁ, Nakashima K, Colak D, Götz M, Fariñas I, Gage FH. Signaling through BMPR-IA Regulates Quiescence and Long-Term Activity of Neural Stem Cells in the Adult Hippocampus. Cell Stem Cell 2010;7:78–89.

38. Martynoga B, Mateo JL, Zhou B, Andersen J, Achimastou A, Urbán N, Berg D van den, Georgopoulou D, Hadjur S, Wittbrodt J, Ettwiller L, Piper M, Gronostajski RM, Guillemot F. Epigenomic enhancer annotation reveals a key role for NFIX in neural stem cell quiescence. Genes Dev. 2013;27:1769–1786.

39. Knobloch M, Pilz G-A, Ghesquière B, Kovacs WJ, Wegleiter T, Moore DL, Hruzova M, Zamboni N, Carmeliet P, Jessberger S. A Fatty Acid Oxidation-Dependent Metabolic Shift Regulates Adult Neural Stem Cell Activity. Cell Reports 2017;20:2144–2155.

40. Leeman DS, Hebestreit K, Ruetz T, Webb AE, McKay A, Pollina EA, Dulken BW, Zhao X, Yeo RW, Ho TT, Mahmoudi S, Devarajan K, Passegué E, Rando TA, Frydman J, Brunet A. Lysosome activation clears aggregates and enhances quiescent neural stem cell activation during aging. Science 2018;359:1277–1283.

41. Morrow CS, Porter TJ, Xu N, Arndt ZP, Ako-Asare K, Heo HJ, Thompson EAN, Moore DL. Vimentin Coordinates Protein Turnover at the Aggresome during Neural Stem Cell Quiescence Exit. Cell Stem Cell 2020;26:558-568.e9.

42. Bechtold B. Violin Plots for Matlab, Github Project. 2016. Available at: https://github.com/bastibe/Violinplot-Matlab.

43. Randi EB, Zuhra K, Pecze L, Panagaki T, Szabo C. Physiological concentrations of cyanide stimulate mitochondrial Complex IV and enhance cellular bioenergetics. Proceedings of the National Academy of Sciences 2021;118:e2026245118.

44. Heaster TM, Walsh AJ, Zhao Y, Hiebert SW, Skala MC. Autofluorescence imaging identifies tumor cell-cycle status on a single-cell level. Journal of Biophotonics 2018;11:e201600276.

45. Chang C-H, Curtis JD, Maggi LB, Faubert B, Villarino AV, O’Sullivan D, Huang SC-C, van der Windt GJW, Blagih J, Qiu J, Weber JD, Pearce EJ, Jones RG, Pearce EL. Posttranscriptional Control of T Cell Effector Function by Aerobic Glycolysis. Cell 2013;153:1239–1251.

46. Walsh AJ, Mueller KP, Tweed K, Jones I, Walsh CM, Piscopo NJ, Niemi NM, Pagliarini DJ, Saha K, Skala MC. Classification of T-cell activation via autofluorescence lifetime imaging. Nat Biomed Eng 2021;5:77–88.

47. Samimi K, Guzman EC, Trier SM, Pham DL, Qian T, Skala MC. Time-domain single photon-excited autofluorescence lifetime for label-free detection of T cell activation. Opt. Lett., OL 2021;46:2168–2171.

48. Shirshin EA, Yakimov BP, Rodionov SA, Omelyanenko NP, Priezzhev AV, Fadeev VV, Lademann J, Darvin ME. Formation of hemoglobin photoproduct is responsible for two-photon and single photon-excited fluorescence of red blood cells. Laser Phys. Lett. 2018;15:075604.

49. Carter P, Smith L, Ryan M. Identification and validation of cell surface antigens for antibody targeting in oncology. Endocrine-Related Cancer 2004;11:659–687.

50. Haustrate A, Hantute-Ghesquier A, Prevarskaya N, Lehen’kyi V. Monoclonal Antibodies Targeting Ion Channels and Their Therapeutic Potential. Front. Pharmacol. 2019;10:606.

51. Fu Z, Zhou J, Chen R, Jin Y, Ni T, Qian L, Xiao C. Cluster of differentiation 19 chimeric antigen receptor T-cell therapy in pediatric acute lymphoblastic leukemia (Review). Oncology Letters 2020;20:1–1.

52. Stringer C, Wang T, Michaelos M, Pachitariu M. Cellpose: a generalist algorithm for cellular segmentation. Nat Methods 2021;18:100–106.

53. Guzman EC, Rehani PR, Skala MC. Cell analysis tools: an open-source library for single-cell analysis of multi-dimensional microscopy images. In: Imaging, Manipulation, and Analysis of Biomolecules, Cells, and Tissues XXI. Vol 12383. SPIE; 2023. p 82–86.

